# Nuclear biocondensate-forming lncRNA and cellular stress surveillance shape host cancer susceptibility

**DOI:** 10.1101/2021.04.26.441543

**Authors:** Anjali Bajpai, Quazi Taushif Ahmad, Sushmita Kundu, Ravi Kant Pandey, Alan Goodman, Apul Goel, Bushra Ateeq, Subhash C. Lakhotia, Pradip Sinha

## Abstract

Host genetics is known to influence cancer susceptibility. However, the specific candidate genes and molecular mechanisms that confer resistance remain poorly understood. Here, we demonstrate the power of haploinsufficiency screen to uncover host genetic regulators of cancer in *Drosophila* and identify the long noncoding RNA (lncRNA) *hsrω*, a structural component of nuclear biomolecular condensate known as omega speckles, as a key host cancer susceptibility locus. Loss of *hsrω* disrupts proteostasis and cell fitness, while its haploinsufficiency accelerates epithelial tumor progression driven by loss of the Lethal giant larvae (Lgl) tumor suppressor. Further validating the breadth of this screening strategy, we independently identified *Drosophila STING* (innate immunity) and *Keap1* (oxidative stress defense) as genetic modifiers of cancer. Moreover, in humans, copy number variations (CNV) in these genes and *Sat III* (a functional human homolog of *hsrω*) correlate with poor cancer prognosis, thereby revealing conserved stress pathways as potential host genetic susceptibility regulators.

## Introduction

Genetic modifiers of cancer are genes that influence cancer risk by interacting with primary driver mutations or oncogenic processes without independently causing cancer. One of the central paradoxes of cancer biology is that, while a variety of oncogenes and tumor suppressors have been identified based on their causal underpinnings of cancer, modifier genes regulating host genetic susceptibility in humans remain underexplored (Castro-Giner et al., 2015; Turnbull et al., 2018). Among the known examples of host genetic modifiers identified by Genome-wide association studies (GWAS), notable are DNA repair genes for oxidative damage (*MUTYH* and *OGG1*, (Raetz and David, 2019), immune checkpoint genes (*CTLA4* and *PDCD1*, (Behrouzfar et al., 2021), or those affecting apoptosis pathways (*BIRC5*, (Siragusa et al., 2024). However, GWAS have limited capacity to identify functional cancer modifiers, which often require validation via haploinsufficiency or heterozygosity tests (Chenevix-Trench et al., 2007; Zeng et al., 2019). In *Drosophila*, genetic tractability enables a rapid assessment of such modifiers (Datta et al., 2023). A classic precedent is the *Minute* (Morata and Ripoll, 1975) class of ribosomal genes (Kongsuwan et al., 1985). Their haploinsufficiency compromises cellular proteostasis (Baumgartner et al., 2021) and disrupts tumor surveillance. Consequently, *Minute* +/- cells are outcompeted by fitter wild type cells or, conversely, fail to compete against tumors, thereby enhancing progression when tumor suppressors are lost (Agrawal et al., 1995; Froldi et al., 2010; Khan et al., 2013).

An emerging theme in cancer biology centres on the role of phase-separated biomolecular condensates, the dynamic cellular entities that regulate processes ranging from signalling and transcription to proteostasis and immune surveillance (Mehta and Zhang, 2022; Shin and Brangwynne, 2017). Disruptions of condensate integrity are increasingly linked to diverse pathologies including tumorigenesis (Banani et al., 2017; Boija et al., 2021). For instance, oncogene activation often utilizes super-enhancer condensates (Sabari et al., 2018) while the loss of PML condensates, for example, compromises p53 availability (Zhang et al., 2024).

In *Drosophila*, the long noncoding RNA-encoding *hsrω* gene codes for multiple transcripts ((Bendena et al., 1989; Lakhotia, 2011), also see **Figure 1A**), the largest nuclear-retained transcript, forms a scaffold for nucleoplasmic condensates known as omega speckles ((Prasanth et al., 2000), also see **Figure 1B**). These condensates dynamically sequester diverse heterogenous ribonuclear proteins (hnRNPs) and RNA-binding proteins (RBPs) to buffer cellular stress (Singh and Lakhotia, 2015). Crucially, loss of *hsrω* transcripts or the HRB87F hnRNP disrupts these condensates (Singh and Lakhotia, 2012). This biology is conserved since the functional human homolog, *Satellite III* (*Sat III*), similarly forms phase-separated nuclear stress bodies dependent on Heat shock factor 1 (HSF1) and sequester RNA-binding proteins (Goenka et al., 2016; Jolly et al., 2002; Jolly et al., 2004). Given the critical role of *hsrω* in stress buffering, we hypothesized that genetic perturbation of this lncRNA would profoundly impact cancer progression.

**Figure 1.**
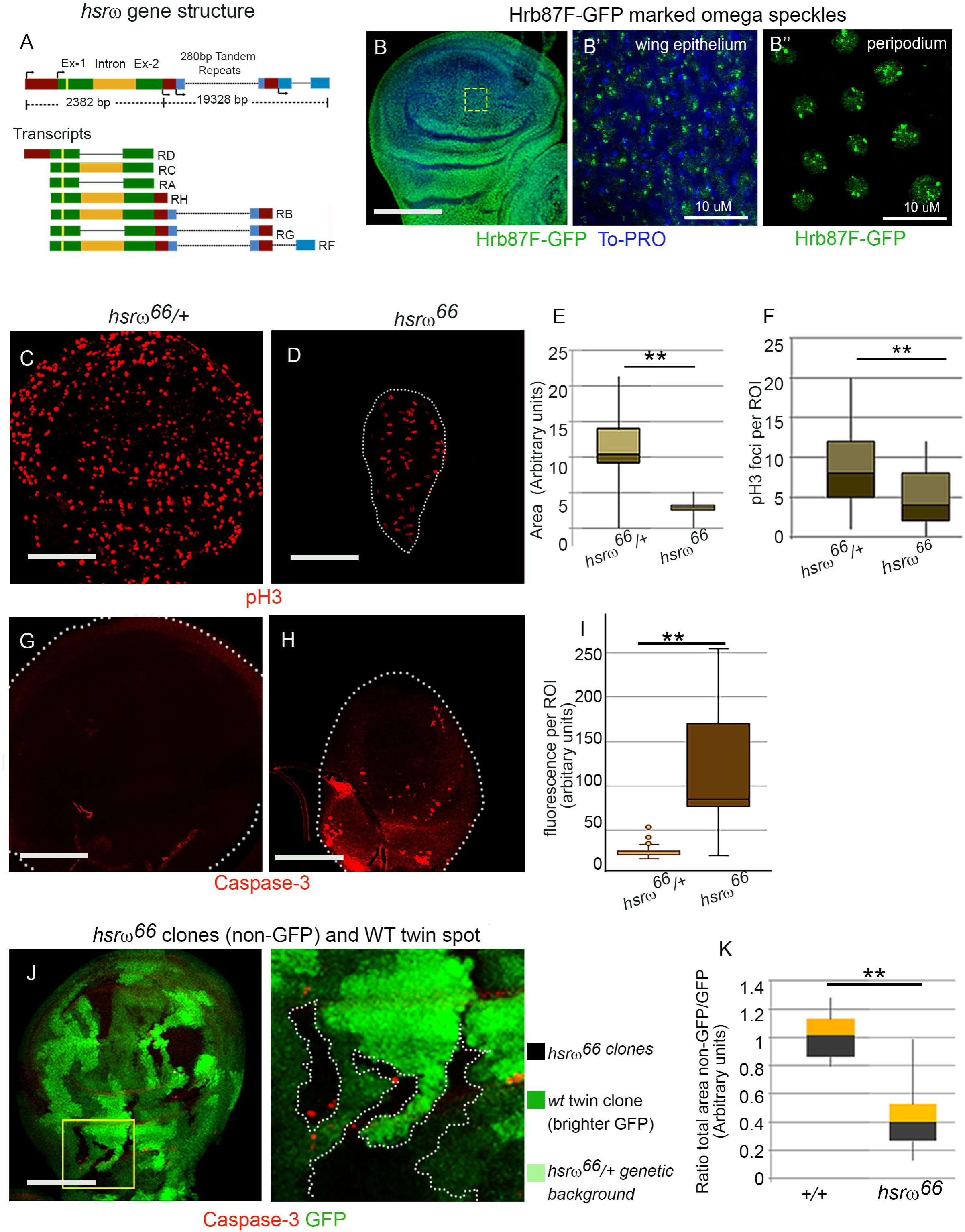
*hsrω* mutant imaginal epithelia display cell stress and undergrowth. (**A**) Diagrammatic depiction of *hsrω* gene and its multiple nuclear and cytoplasmic transcripts (Sahu et al., 2020). The larger nuclear transcripts (RB, RG and RF) carry a long stretch of tandem repeats, providing a physical scaffold for binding hnRNPs, such as Hrb87F (Prasanth et al., 2000). (**B**) Nuclear omega speckle biocondensates, marked by GFP-tagged Hrb87F (green) in wing imaginal disc cells. Nucleus is marked by To-PRO (blue). (**C–F**) Third instar wing imaginal discs of *hsrω*^66^/+ (C) and *hsrω^66^* (D) larvae immunostained for phospho-Histone 3 (pH3, red) to mark dividing cells, and quantification for wing disc size (E) and relative abundance of pH3-positive nuclei per arbitrary unit area (F). (**G–I**) Expression of Caspase-3 (red) in *hsrω*^66^/+ (G) and *hsrω^66^* (H) wing discs and quantification of Caspase-3 (fluorescence per ROI, I) (**J-K**) Mosaic wing disc with *hsrω^66^* somatic clones (non-GFP) and their *hsrω^66^*/+ twins (bright green, due to two copies of GFP). Boxed area in J is shown at a higher magnification to reveal Caspase-3 expression (red) in *hsrω^66^* cells abutting the clone’s border. Comparison of *hsrω^66^*clone areas with the twin-spot regions (K). Scale bars 100µm in all confocal images, except in B’ which is 10 µm.

Here, we employed a hypothesis-driven candidate gene approach to uncover host genetic modifiers of cancer. We demonstrate that the omega speckle-associated lncRNA, *hsrω*, acts as a potent host genetic modifier of cancer susceptibility by regulating proteostasis. Furthermore, to validate the breadth of this haploinsufficiency screening approach, we expanded our search to other arms of the cellular stress response. We demonstrate that *Drosophila STING* (*dSTING*), a regulator of innate immune stress (Cai et al., 2020; Martin et al., 2018) and *Keap1*, a regulator of oxidative stress (Sykiotis and Bohmann, 2008) also serve as independent host genetic modifiers. Collectively, our findings highlight a conserved network of stress surveillance pathways that define tumor susceptibility.

## Results

### Study design

To systematically uncover how host genetics contributes to intrinsic epithelial tumor surveillance, we employed a candidate gene approach focusing on haploinsufficiency. We selected distinct but potentially overlapping stress regulators: the nuclear biomolecular condensate-associated lncRNA, *hsrω*, the innate immunity pathway component, *dSTING* and the redox regulator Keap1. Our strategy involved a stepwise characterization: first, we defined the consequences of *hsrω* loss on tissue growth, proteostasis, and cellular fitness in normal epithelium. Next, we tested whether *hsrω* haploinsufficiency facilitates the neoplastic transformation of somatic clones lacking the tumor suppressor *Lgl*. Finally, extending the logic of stress surveillance, we examined whether haploinsufficiency of *dSTING*—a key sentinel of the cGAS–STING pathway—mirrors the tumor-promoting effects of *hsrω* loss, thereby establishing a broader role for stress regulators in host cancer susceptibility.

### *hsrω* mutants are growth-compromised and display disrupted proteostasis

Homozygosis for the *hsrω* near-null mutant, *hsrω^66^*(Johnson et al., 2009), causes high embryonic lethality, with the escapers emerging as weak short-lived adults . Analysis of these larvae revealed undergrown wing imaginal discs with significantly fewer dividing cells per unit area than controls (**Figure 1C–F**) and elevated cell death (**Figure 1G, H**). The *hsrω^66^* somatic clones were similarly smaller than their wild type twin spots, with cell death evident along clone borders (**Figure 1I, J**)—a phenotype characteristic of ‘loser’ cell behaviour in cell competition (de la Cova et al., 2004; Moreno et al., 2002). These growth defects prompted us to examine whether loss of *hsrω* perturbs the protein homeostasis landscape in developing epithelial cells.

Proteomic profiling revealed a collapse in the translation machinery. Comparing the proteome of mutant and wild type (WT) larval wing epithelium, we noted >±2-fold (p<0.05) changes in 73 proteins in *hsrω^66^* mutants (**Figures 2A and S1A, B**). We observed a specific depletion of ribosomal structural proteins (RpS15Ab, mRpL50, and mRPL24; **Figure 2B**) alongside proteins involved in protein stability such as the heat shock protein Hsp23 and protein turnover, the SUMO-protease Ulp1 (**Figure S1C**). Rank-based enrichment analysis confirmed the overall downregulation of cytosolic and mitochondrial large ribosomal subunits (**Figure 2C**), as well as the cytoplasmic and mitochondrial translation machinery (**Figure 2D**). Furthermore, RNA Pol II transcription machinery components and the mediator complex were depleted (**Figures S1E**), indicating a systemic failure in gene expression and protein synthesis control. We noted that the *hsrω^66^* proteome showed an upregulation of stress response markers (**Figure S1D**), including the cytoplasmic Glutathione S-transferase E6 (GstE6) (Aloke et al., 2024) and mitochondria-associated Mgstl (Toba and Aigaki, 2000), indicative of redox imbalance. Such cellular changes likely prefigure DNA damage signatures, consistent with the observed elevation of Rpa1, a DNA double-strand break-associated protein (**Figures 2E**).

**Figure 2.**
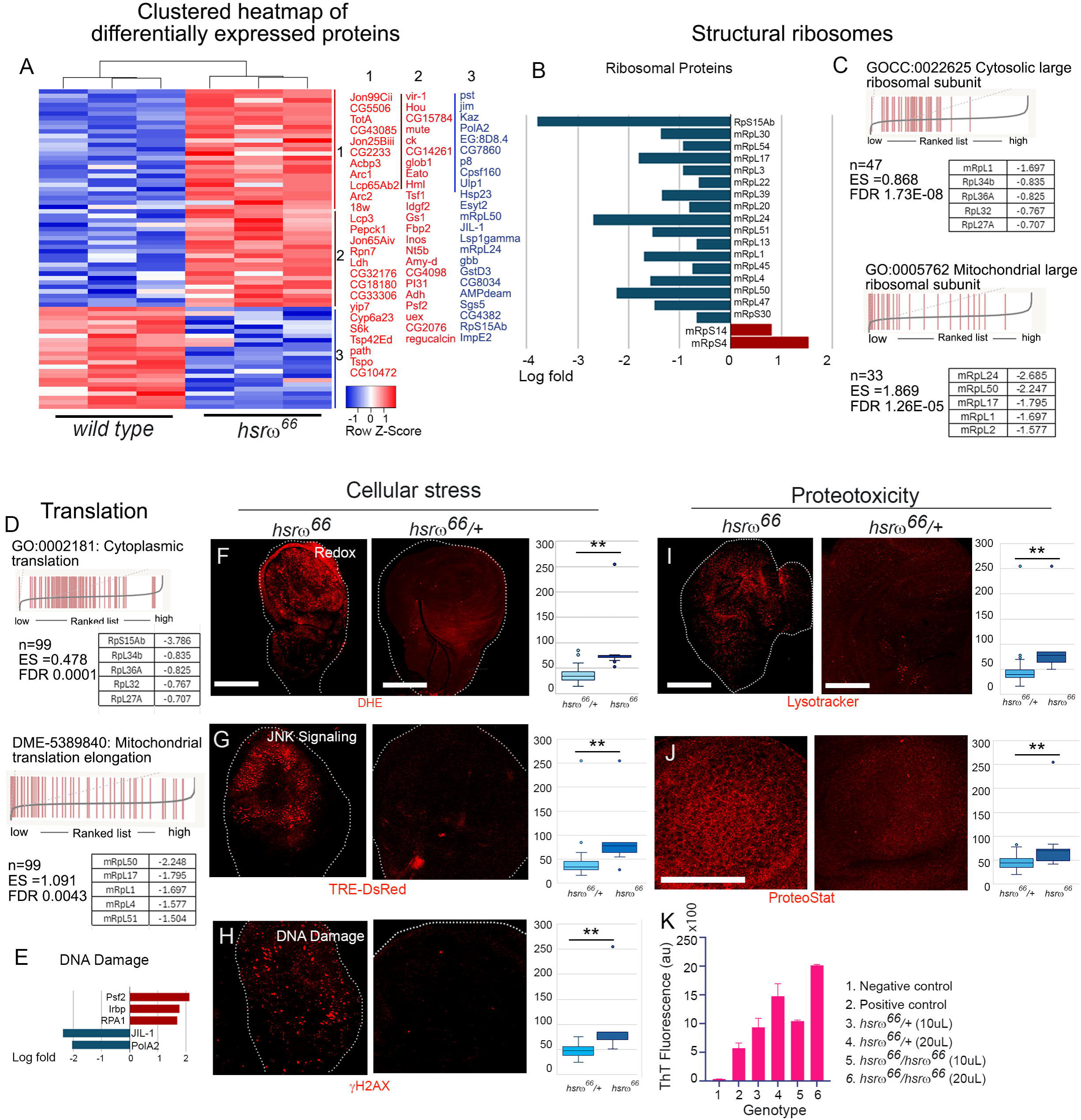
*hsrω^66^* epithelium reveals cellular and proteotoxic stress. (**A**) Clustered heatmap displaying status of proteins significantly altered (±>log_2_ 2-fold, p < 0.05, q<0.05) in *hsrω^66^* versus wild type wing imaginal disc proteome. On the right is the list of proteins upregulated (red) or downregulated (blue) in the *hsrω^66^* proteome. (**B**) Relative abundance (log_2_ fold change) of select ribosomal structural proteins in *hsrω^66^* proteome compared to its wild type counterpart. (**C-D**) Rank-based Gene Ontology (GO) enrichment analysis of cytosolic and mitochondrial ribosome proteins (C) and translation machinery (D). The x-axis represents the list of proteins in ascending order of fold change in *hsrω^66^* versus wild type. Black bars along the x-axis mark the position of proteins being queried. The tables alongside give the protein levels of the top five mis-expressed proteins belonging to a given GO class. **(E**) Relative abundance of key DNA damage-associated proteins in *hsrω^66^* proteome. (**F-H**) Representative examples of wing imaginal discs displaying signatures of cellular stress: ROS seen using superoxide-sensitive, Dihydroethidium (DHE, F); JNK signalling visualized using a bio-sensor, TRE-dsRED, for JNK-mediated AP-1 signalling (G); DNA double-strand breaks, marked by γH2AX (H), in *hsrω^66^* and *hsrω^66^*/+ counterpart and its quantification on the right. The Y-axis of the graphs represents the fluorescence per ROI. (I-K) Signatures of increased proteotoxicity in *hsrω^66^* versus *hsrω^66^*/+ wing epithelium: Elevated LysoTracker fluorescence (I) and protein aggregates visualized using fluorescent dye ProteoStat (J) in *hsrω^66^* and *hsrω^66^* /+ wing imaginal discs. Quantification of Thioflavin fluorescence in cell homogenates of wing discs of *hsrω^66^* and control larvae a measure of protein aggregates. Two-tailed Student’s t-test was applied to determine the statistical significance between test and controls in 2F-J; *P≤ 0.05 and **P≤ 0.001. Error bars represent mean ± SEM. ES: Enrichment Score. p-value is the Nominal p-value. A negative value of NES indicates poor enrichment of the queried geneset in *hsrω^66^* mutant proteome versus control proteome. Scale bars: 100 µm.

We further validated these proteomic signatures using cellular stress markers. The *hsrω^66^* mutant wing epithelium displayed elevated ROS levels (DHE staining; **Figure 2F**) and enhanced JNK signaling (**Figure 2G**). This oxidative stress was accompanied by augmented γH2AX staining, confirming the presence of DNA double-strand breaks (Mah et al., 2010 ; **Figures 2H**). We reasoned that a suboptimal proteome under such stress would lead to the accumulation of damaged proteins, necessitating increased lysosomal clearance (Settembre et al., 2013). In agreement, *hsrω^66^* wing discs showed increased acidic lysosomes detected by LysoTracker (**Figure 2I**) and an increase in Ubiquitin-positive foci (**Figure S2G**) indicating proteins marked for proteasomal degradation (Martínez-Férriz et al., 2022). Finally, we observed signatures of increased proteotoxicity in the wing epithelium owing to loss of proteostasis. For instance, the mutant proteome showed a marked elevation of ARC1 and ARC2 (**Figure 2A**), proteins associated with pathological aggregates (Schulz et al., 2023). Crucially, *hsrω^66^* wing epithelium exhibited marked increase in cellular uptake of ProteoStat (**Figure 2J**), a dye that fluoresces upon intercalation with protein aggregates (Baumgartner et al., 2021), and a pronounced increase in Thioflavin fluorescence in cell homogenates (**Figure 2K**), both telltale markers of elevated proteotoxicity (Kandel et al., 2024).

These findings indicate that *hsrω* loss disrupts ribosome integrity and proteome surveillance, fostering oxidative stress and proteotoxicity. The resulting translational stress therefore compromises cellular fitness, predisposing *hsrω^66^* cells to elimination (**Figure 1I**) through competition with the neighboring wild type cells. Thus, the cell-competition phenotype of *hsrω* mutants—and, by extension, the functional compromise of the omega speckle biomolecular condensate—culminates in impaired cellular proteostasis and compromised stress adaptation.

### *hsrω* displays haploinsufficiency for tumor suppression

We next tested the survival of *lgl* somatic clones in *hsrω^66^/+* larvae, given that perturbed proteostasis permits tumor progression in other contexts, such as the *Minute* genetic background (Baumgartner et al., 2021; Khan et al., 2013). In a wild type epithelium, somatic clones lacking Lgl are recognized and eliminated by a tissue surveillance mechanism known as cell competition (Khan et al., 2013; Menéndez et al., 2010; Suijkerbuijk et al., 2016). These clones display increased cellular stress (**Figure S2**), exhibit elevated cell death, and are basally extruded ((Khan et al., 2013), also see **Figure 3A**). By contrast, *lgl* clones generated in *hsrω^66^/+* wing imaginal discs evaded elimination and underwent neoplastic transformation, marked by disrupted cortical F-Actin (**Figure 3B, C**). Consistent with the acquisition of a ‘winner’ cell state, we observed cell death in *lgl^+^/lgl* cells along the *lgl* clone boundaries (**Figure 3B**). Furthermore, these transformed *lgl* clones displayed elevated cell proliferation (pH3, **Figure 3D**) and an increase in Matrix Metalloproteinase-1 (MMP1; **Figure 3E**), indicating ECM degradation and the acquisition of invasive properties (Beaucher et al., 2007). We thus hypothesized that the compromised proteostasis in the *hsrω* mutant background (**Figure 2**) would render the epithelium hypersensitive to proteosome inhibitor. Indeed, administration of the classical proteasome inhibitor MG-132 (Chen et al., 2011) compromised the tumor susceptibility of *hsrω^66^/+* larvae, reducing *lgl* tumor burden in a dose-dependent manner (**Figure 3F**). This suggests that the tumor-promoting environment of *hsrω* haploinsufficiency relies on a precarious proteostasis balance.

**Figure 3.**
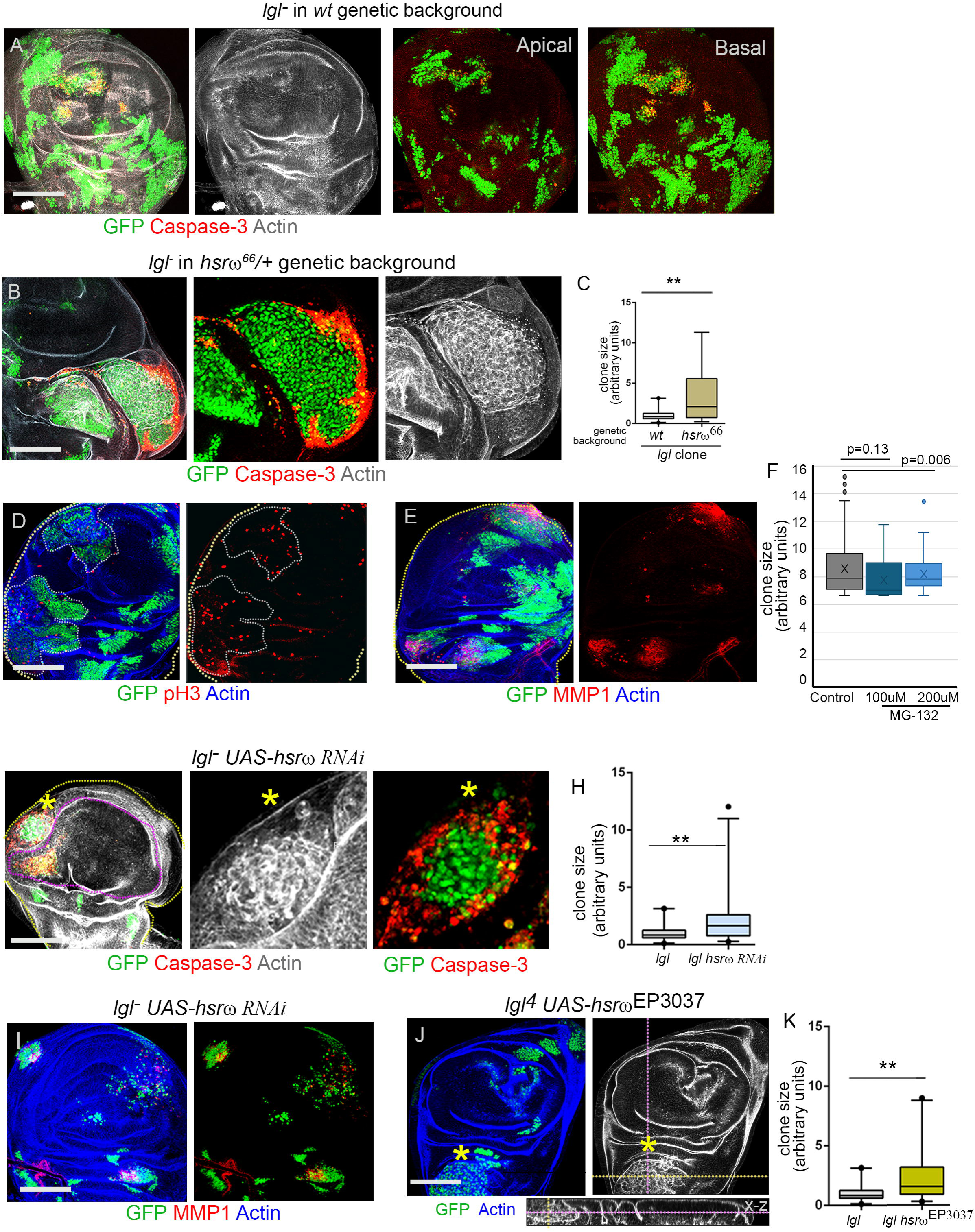
Haploinsufficiency of *hsrω* is a tumor-permissive genetic background. (**A**) GFP-marked *lgl^−^* somatic clones (GFP, green) induced using heat shock flippase, generated in a wild type genetic background. *lgl* clones (*hsflp; lgl^4^FRT40*) display cell death (Caspase-3, red). Note enrichment of Caspase-positive *lgl* clones in basal compared to apical projections of select optical sections on the right panel. (**B**-**C**) *lgl^−^* somatic clones (GFP, green) in a heterozygous *hsrω^66^/+* background (*hsflp; lgl^4^FRT40; hsrω^66^/+*) undergo neoplastic transformation revealed by disrupted F-Actin. Note the characteristic large, dispersed nuclei (GFP) of transformed clones, and Caspase-3 expression along boundaries. Quantification of *lgl* clone area in wild type (WT) vs. *hsrω^66^/+* wing imaginal discs (C). (**D-E**) *lgl^−^* clones in *hsrω^66^/+* imaginal discs displaying excessive proliferation (pH3, D) and elevated levels of Matrix Metalloproteinase (MMP1) expression (E). (**F**) Graph displaying reduction in clone size of *lgl; hsrω^66^/+* tumors, when larvae were fed proteosome inhibitor MG-132 at two different concentrations. (**G-I**) *lgl* somatic clones (marked in GFP) with co-down regulation *(hsflp; lgl^4^ UAS-hsrω-RNAi FRT40A* (G-I) or up regulation (*lgl^4^ FRT40A UAS-hsrω^EP3039^*; J) of in *WT* genetic background. *lgl^−^*clones survive and undergo neoplastic transformation (stars), marked by disrupted F-Actin (Phalloidin staining) and induce exaggerated Caspase-3 expression along the clone boundary (G). Quantification of *lgl^−^* clone areas with loss (H) or gain (K) of *hsrω*. Two-tailed Student’s t-test was applied to determine the statistical significance between test and controls in 3C, F. H and K; *P≤ 0.05 and **P≤ 0.001. Scale bars: 100 µm.

We next asked if this tumor susceptibility was driven by the loss of structural integrity of the omega speckle. A hallmark of condensate-based regulatory systems is their sensitivity to component dosage—both depletion and overexpression can perturb phase separation (Shin and Brangwynne, 2017) also recapitulated by omega speckles that are sensitive to doses of *hsrω* and omega speckles-associated hnRNPs such as Hrb87F (Lakhotia, 2011; Prasanth et al., 2000). Consistent with this model, we found that *lgl* somatic clones underwent neoplastic transformation not only upon *hsrω* knockdown (*UAS-hsrω-RNAi*; **Figure 3G-I**) but also upon overexpression (*UAS-hsrω^EP3037^*; **Figure 3J, K**). Under both conditions, *lgl* clones gained a growth advantage, induced cell death in the neighboring cells (**Figure 3G**), and upregulated invasion markers (MMP1; **Figure 3I**).

To confirm that condensate integrity is the critical factor, we manipulated other structural components. Heterozygosity of *Df(3R)Hrb87F*—which encodes an hnRNP localized to omega speckles—similarly promoted *lgl* transformation (**Figure S3A**). Moreover, downregulating the chromatin remodeling ATPase *ISWI*—required for omega speckle structural integrity (Onorati et al., 2011; Singh and Lakhotia, 2015)—also facilitated neoplastic transformation (**Figure S3C**). Finally, *lgl* somatic clones induced in the adult intestinal epithelium of *hsrω^66^/+* adult flies also displayed neoplastic transformation (**Figure S4**), unlike those generated in a WT background. This indicates that *hsrω* haploinsufficiency creates a tumor-susceptible host genetic background, likely through sensitivity to biocondensates disruption and ensuing impaired protein homeostasis.

### Tumor inhibitory role of redox regulators Nrf2-Keap1 and inflammatory protein dSTING

To validate the versatility of haploinsufficiency screening in uncovering diverse host genetic determinants, we extended our analysis to two other functional classes of stress regulators identified in our screen: the redox sensor Nrf2-Keap1 and the viral immune sensor cGAS-STING. While somatic variations in these pathways are suspected to influence cancer risk (Michalak and Michalak, 2025) their direct impact on intrinsic tumor surveillance remains to be functionally dissected in a host genetic model.

First, we examined the Nrf2-Keap1 oxidative stress pathway. This conserved system mediates cellular responses to oxidative damage as Keap1 suppresses the transcription factor Nrf2 (*Drosophila* homolog, CncC) by targeting it for proteasomal degradation (Fuse and Kobayashi, 2017). In the absence of Keap1, CncC translocates to the nucleus to activate antioxidant and proteostasis genes (Radhakrishnan et al., 2010). We reasoned that altering this redox balance might shield emerging tumor cells from oxidative stress-induced elimination. Consistent with this, *lgl* somatic clones generated in a *Keap1* haploinsufficient background (*Keap1^EY5^/+*) exhibited enhanced growth and neoplastic transformation (**Figure 4A**) when compared to *lgl* clones generated in wild type genetic background. We observed a similar tumor-promoting effect upon cell-autonomous downregulation of *Keap1* (*lgl; UAS-Keap1-RNAi*; **Figure 4B, C**) or direct genetic gain of *CncC* (*lgl; UAS-CncC*; **Figure 4D**), with transformed *lgl* clones displaying elevated cell death in abutting wild type neighbors (**Figure 4C**). This suggests that constitutive activation of antioxidant defense (via *Keap1* loss) acts as a barrier to the elimination of oncogenic clones.

**Figure 4.**
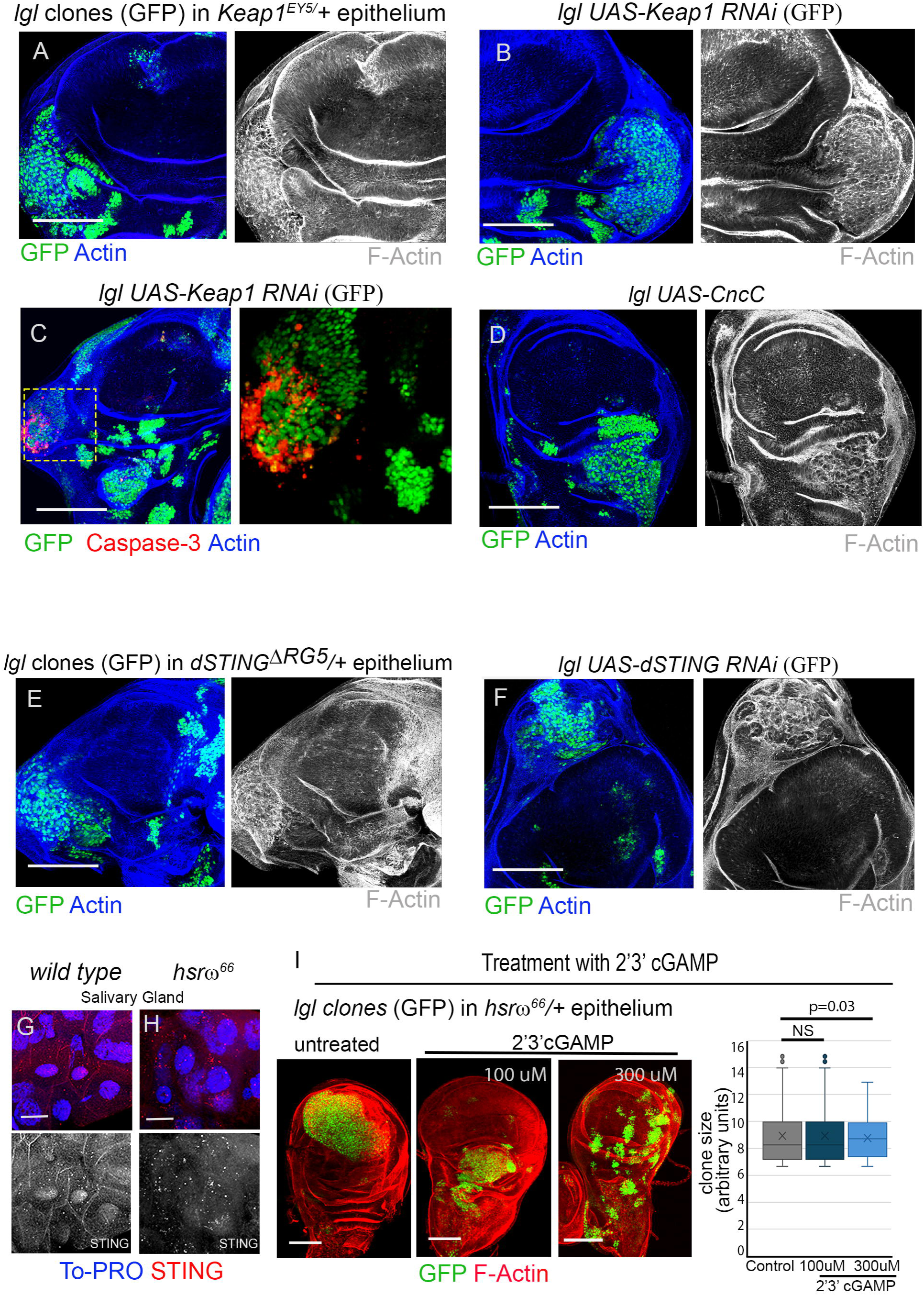
Redox regulator *Keap1/Nrf2* and inflammatory gene *dSTING* are haploinsufficient for tumor suppression. **(A-D)** GFP-marked *lgl* somatic clones in a *Keap1* heterozygous genetic background (*lgl^4^; Keap^EY5^/+*, A), or upon cell autonomous downregulation of *Keap1* (*lgl^4^; UAS-Keap1 RNAi*, B), or gain of downstream regulator CncC (*lgl^4^; UAS-CncC,* C), display neoplastic transformation (disrupted F-Actin) and cell death along clone boundary (Caspase-3, C). **(E-F)** GFP-marked *lgl* somatic clones in dSTING heterozygous (*lgl^4^; STING^ΔRG5^/+*, E) or STING down regulated (*lgl^4^; UAS-dSTING RNAi*, F) genetic background display neoplastic transformation with disrupted F-Actin. **(G-I)** Disrupted sub-cellular distribution of dSTING in cells of *hsrω^66^* mutant salivary gland (H) with enhanced punctae and reduced general cytoplasmic distribution compared with a uniform distribution and fewer punctae in wild type gland (G). (I) Representative wing discs from larvae bearing *lgl^−^* somatic tumors generated in *hsrω^66^/+* genetic background and fed on STING agonist, 2′3′-cGAMP, at two different concentrations; quantification of *lgl* clone size from unfed and fed larvae (far right). Two-tailed Student’s t-test was applied to determine the statistical significance between test and controls in 4I. NS: Not Significant.

Next, we investigated the cGAS–STING pathway. This pathway acts as a sentinel for aberrant cytosolic DNA—a hallmark of genomic instability and cellular damage or of virus-induced biotic stress (Hopfner and Hornung, 2020). In *Drosophila*, activation of dSTING triggers an inflammatory response via the NF-κB/Relish pathway (Cai et al., 2020; Martin et al., 2018). We hypothesized that haploinsufficiency of *dSTING* would impair the host’s ability to sense and eliminate early oncogenic clones. Indeed, *lgl* somatic clones generated in a *dSTING* mutant background (*dSTING^ΔRG5^/+*) showed enhanced survival and underwent neoplastic transformation (**Figure 4E**), effectively phenocopying the *hsrω^66^/+* tissue. Similarly, cell-autonomous *dSTING* knockdown within *lgl*-deficient clones led to robust tumor formation (**Figure 4F**), confirming that innate immune sensing is required for effective tumor surveillance.

Finally, we explored the potential crosstalk between these mechanisms. While identified independently, our data suggests a functional link between nuclear condensates and immune surveillance. Notably, we observed that compared to wild type larval salivary glands, the *hsrω^66^* mutant cells displayed disrupted dSTING punctae with reduced diffuse distribution in cytoplasm (**Figure 4G, H** and **Figure S5**). This suggests that the condensate defects in *hsrω* mutants may compromise dSTING trafficking or assembly. This was confirmed by marked suppression of *lgl* tumor growth following treatment with the dSTING agonist (**Figure 4I**) 2′3′-cGAMP (Cai et al., 2020). This demonstrates that pharmacological reactivation of innate immunity can compensate for the surveillance deficits caused by *hsrω* loss, highlighting a functional intersection between nuclear condensates and immune signaling.

### CNVs in human homologs of stress regulators and Sat III correlate with cancer severity

To evaluate the clinical relevance of our findings in *Drosophila*, we investigated whether genetic alterations in the human homologs of these stress regulators correlate with patient outcomes. First, we examined if variations in gene dosage affect the survival of cancer patients using the Cancer Genome Atlas (TCGA). We analysed Copy Number Variations (CNVs) in human *KEAP1* and *STING1* across diverse cancer types (**Figures S6A, B)**. Although TCGA data primarily reflects somatic alterations of tumor origin, it provides a window into how gene dosage impacts progression. Kaplan-Meier survival plots revealed a significant correlation between CNVs in these genes (including deletions and amplifications) and poor patient survival (**Figures S6A, B**). This confirms that, similar to our *Drosophila* models, the dosage of these stress regulators is a critical determinant of human cancer prognosis.

Next, we focused on the human functional analog of *hsrω*, the *Satellite III* (*Sat III*) repeats. While *hsrω* lacks a sequence homolog in mammals, the *Sat III* repeats at the pericentromeric region of chromosome 9 (9q12) are its functional analog (Jolly and Lakhotia, 2006; Jolly et al., 2004). Like *hsrω*, *Sat III* is actively transcribed during stress to form phase-separated nuclear ‘stress bodies’ (Jolly et al., 2004; Goenka et al., 2016). These bodies sequester Heat Shock Factor 1 (HSF1) and specific RNA-binding proteins. Since repetitive *Sat III* sequences are not resolved in standard TCGA exome data, we first looked at its obligate binding partner, HSF1. Notably, TCGA analysis revealed that CNVs in HSF1 are also associated with poor patient survival (**Figure 5A, B**), thereby supporting the importance of the stress body machinery. Finally, we tested the hypothesis that germline variations in *Sat III* itself act as a host susceptibility factor. Since *Sat III* resides in a large heterochromatic block of pentameric repeats at 9q12 (**Figure 5C**), we utilized digital droplet PCR (ddPCR) to precisely quantify copy number variations in *Sat III* using the 9q12-specific, pHuR98 probe (de Lima et al., 2021; Jolly et al., 2002). We analysed genomic DNA from prostate cancer patients and compared them to benign controls. Strikingly, we observed significant germline copy number variations in *Sat III* in patients with high cancer severity (Gleason scores 6–10) compared to benign controls (**Figure 5D**). Further, reminiscent of the tumor-promoting effects of *hsrω* haploinsufficiency in flies, we also observed an overall decrease in Sat III transcript levels across different prostate cancer cell lines compared to benign control (**Figure 5E**). This agrees with previous report that *Sat III* is misregulated in cancer cell lines (Chatterjee and Sengupta, 2022).

**Figure 5.**
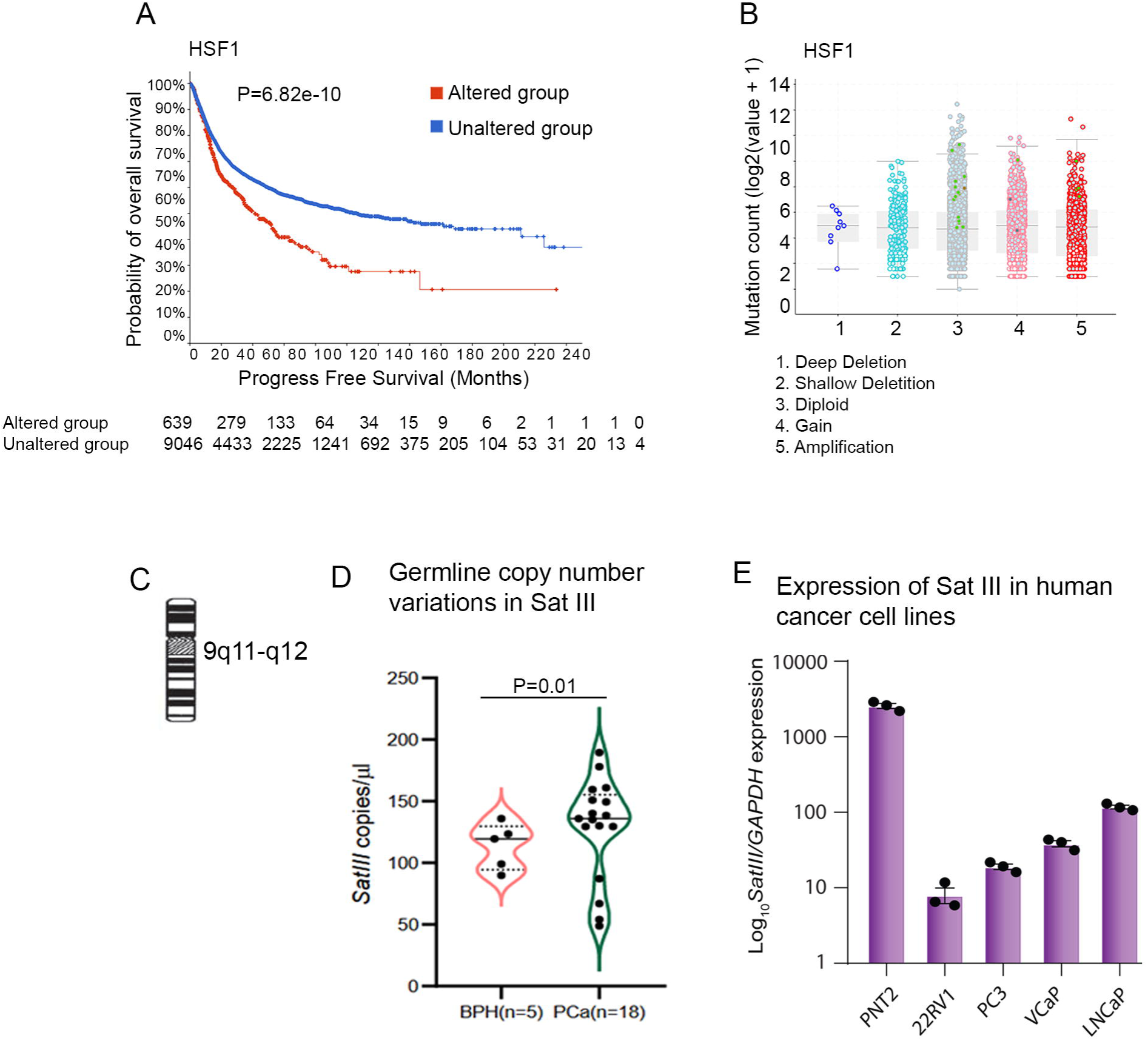
Genetic variations in Sat III in human cancer patients. **(A-B)** Kaplan-Meier curves (generated on cBioPortal platform), comparing survival of individuals with epithelial cancer carrying Copy Number Alterations (CNA) in *HSF1* with individuals not carrying any alteration in *HSF1*. The y-axis represents the probability percent survival, and the x-axis shows progress free survival in months. The plot was generated by querying 91469 samples across 233 studies and included 16 cancer types, (A); Graph displaying the distribution of various genetic alterations in *HSF1* across patients (B). (**C-E**) Schema depicting *Sat III* locus on pericentromeric region of chromosome 9 (C), expression of *Sat III* in benign and prostate cancer cell lines (D). Violin plot depicting germline copy number variations of *Sat III* in PCa patients compared to benign control, BPH (E).

Collectively, our findings in *Drosophila* and human cancers show that perturbations in the dosage of stress surveillance components, whether the protein coding *KEAP1* and *STING1* or the noncoding *hsrω*/*Sat III* architectural RNAs, are broadly associated with cancer susceptibility, severity and poor prognosis.

## Discussion

Our study establishes *Drosophila* haploinsufficiency screen as a powerful tool for identifying host genetic regulators of cancer susceptibility **(Figure 6)**, a class of genes often overlooked by traditional oncogene/tumor suppressor screens. While the roles of high-penetrance mutations (e.g., TP53, PTEN) are well defined (Berger and Pandolfi, 2011), the genetic architecture of host resistance remains underexplored. We demonstrate that subtle genetic deficits (haploinsufficiency) in fundamental homeostatic regulators, specifically the nuclear condensates, redox sensors, and immune sentinels, do not initiate tumors per se, but severely erode the ‘surveillance buffer’ of cells. This renders the epithelium permissive to neoplastic transformation upon a second hit, such as loss of the cell polarity protein Lgl.

**Figure 6.**
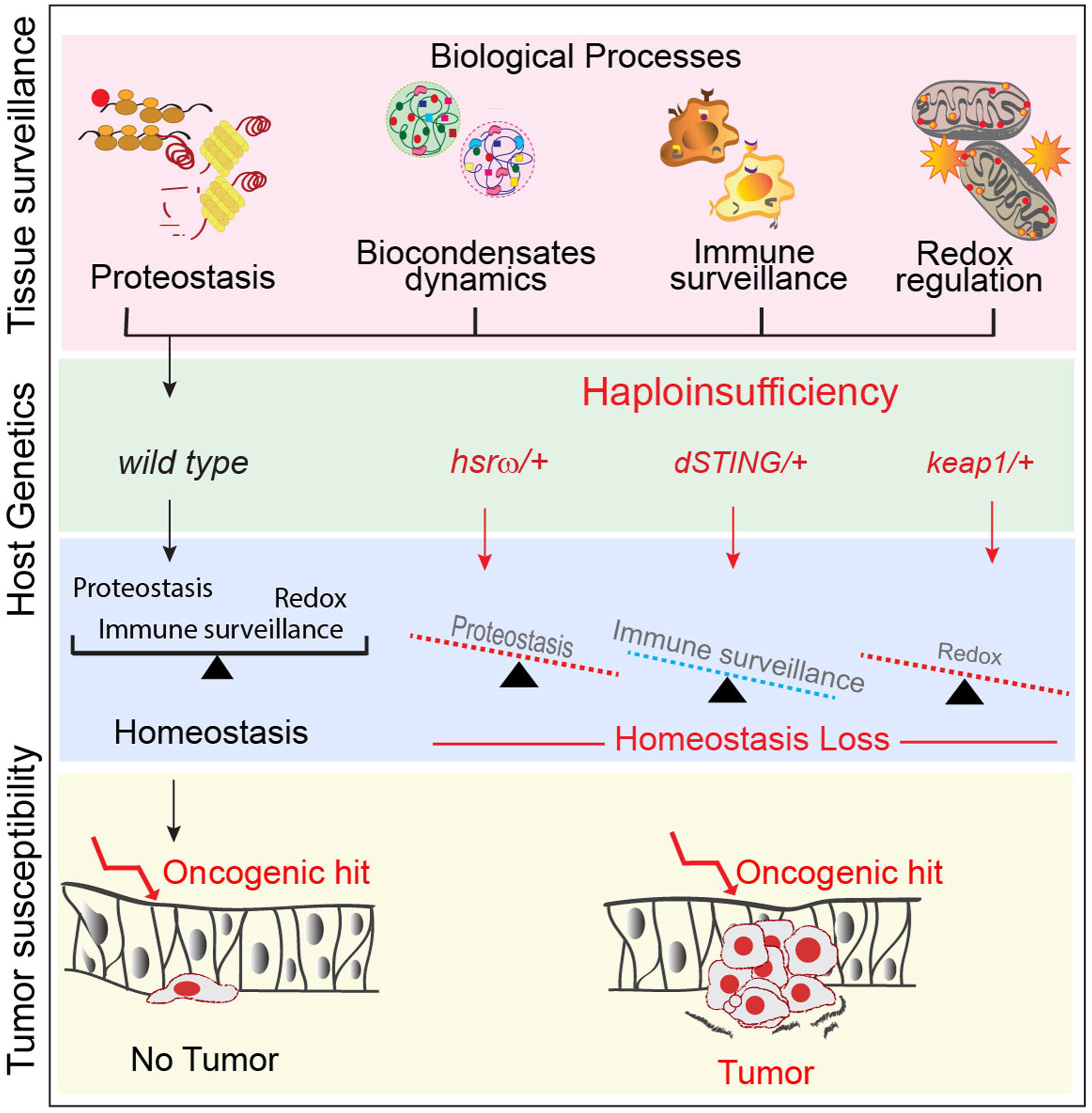
Summary: Schema displaying haploinsufficiency as a strategy to uncover host genetic modifiers of cancer susceptibility. Moderate genetic variations (haploinsufficiency) transform a tumor-resistant wild type epithelium to a genetically altered tumor-permissive epithelium (right panel).

A major finding of this study is the identification of the lncRNA-encoding *hsrω*, essential for formation of omega speckle biomolecular condensates, as a key suppressor of susceptibility. This expands the “Proteostasis Addiction” model of cancer (Dai and Sampson, 2016). We propose that *hsrω*- dependent omega speckle condensates function as proteostasis regulator by, sequestering various RNA-binding proteins, and buffering the translational stress. Loss of this buffer, as in *hsrω* haploinsufficiency, creates a ‘proteostasis-vulnerable’ epithelium that cannot cope with the physiological demands of emerging oncogenic clones, which effectively lowers the threshold for tumorigenesis. This mirrors the classic Minute phenotype in *Drosophila* (Baumgartner et al., 2021), but with a crucial distinction that in this case a non-coding RNA rather than a ribosomal protein acts as the linchpin of surveillance.

Validating the breadth of our screening strategy, we independently identified *STING* (an innate immunity factor) and *Keap1* (a redox regulatory factor) as host modifiers. While distinct in function, these pathways likely converge. Our observations that *hsrω* loss disrupts the cellular distribution of dSTING, and that STING agonists can rescue the *hsrω* tumor susceptibility phenotype suggest a functional intersection. We postulate that the dynamics of omega speckle nuclear condensates also regulate the nucleocytoplasmic distribution or availability of specific immune or redox factors, such as STING. Thus, *hsrω*, *STING*, and *Keap1* may function as nodes in an integrated ‘Stress Surveillance Network.’ Disabling any node, whether by compromising activities of the hsrω or the sensors (STING/Keap1), collapses the network, which in turn permits cells carrying tumorigenic hits overcome tumor suppressive cell competition.

A significant implication of our study is the potential translational link to human *Sat III* repeats. Although the *Sat III* repeats have long been considered “junk DNA” or mere markers of stress, in context of the *Drosophila* model, we suggest a significant tumor-suppressive function to this non-coding locus. Our finding that the germline Copy Number Variations (CNVs) in *Sat III* correlate with prostate cancer severity suggests that the highly polymorphic repetitive regions of the human genome, often ignored in standard exome sequencing, may harbor critical determinants of cancer susceptibility. Just as *hsrω* dosage dictates fly cells’ tumor resistance, the intrinsic copy number of human *Sat III* may determine the robustness of a patient’s “cellular stress response”, thereby influencing clinical prognosis.

Finally, our findings resonate with the evolutionary concept that cancer resistance is a distinct trait from tumor suppression (Klein, 2009). Long-lived mammals, such as elephants or whales, have evolved redundant resistance mechanisms, including increased TP53 copy numbers (Sulak et al., 2016). We propose that *hsrω/Sat III*, STING, and Keap1 represent a similar layer of ‘background’ resistance. These genes do not drive the cancer, but they set the rules of engagement. A host with robust condensate dynamics and immune sensing (high resistance) forces oncogenes to overcome a steep barrier, whereas a host with haploinsufficient defects (low resistance) provides a more permissive environment. Combining *Drosophila* genetics with human CNV analysis opens a new avenue to systematically map these elusive ‘host resistance’ genes, moving the field beyond the tumor-centric driver model towards a more holistic understanding of cancer susceptibility.

## Acknowledgement

We acknowledge Bloomington Drosophila Stock Centre (Indiana University) for fly stocks and Developmental Studies Hybridoma Bank (University of Iowa) for antibodies. Ethical approval for use of patient samples in this study was obtained from the Institutional Ethics Committee (IEC) at Indian Institute of Technology Kanpur (Approval Number: IITK/IEC/2020-21/II/05) issued on February 18, 2021 for five years. All patients participating in this study provided written informed consent. We thank S. Ganesh, Indian Institute of Technology Kanpur, for the gift of the FK2 anti-ubiquitin antibody. This work was supported by the Department of Biotechnology, Govt. of India and Wellcome Trust India Alliance (IA/E/13/1/501271) and Department of Science and Technology, DST/WOS-A/LS-75/2021 to AB. Support by the Science & Engineering Research Board (SERB), Department of Science and Technology, Govt. of India (CO/S5/AB-03/2017) to PS and DBT/ Wellcome Trust India Alliance (IA/S/19/2/504659) to BA is acknowledged. SCL is supported by the Department of Biotechnology, Ministry of Science and Technology (BT/PR32126/BRB/10/1775/2019) and the Science & Engineering Research Board, Govt. of India, as a Distinguished Fellow (SB/DF/009/2019).

The authors declare no competing interests.

## Material and Method

### Fly Stocks

Fly stocks were maintained at 25° C on standard fly food containing corn powder, yeast, and sugar. The following mutant/transgenic lines were used in this study: *w; l(2)gl^4^ FRT40A/CyO* (#36289, BDSC); *w; w; hsr*ω*^66^/TM6B* (Johnson et al., 2011); *w;UAS-hsr*ω-*RNAi(2x)*(Mallik and Lakhotia, 2009); *EP3037/ EP3037* (Mallik and Lakhotia, 2009); *Df(3R)Hrb87F/Tb* (#67701, BDSC); *y w hs-flp, tub-Gal4, UAS-GFP/; tub-Gal80 FRT40A/CyO; w; FRT82B* (#86313, BDSC); *y w hs-flp; FRT82B Ubi-GFP* (#5188, BDSC); *w; TRE-DsRed;+ (*#59011, BDSC);*UAS-STING RNAi* (#31565, BDSC); *w;+;th-lacZ* (Bennett and Harvey, 2006); *STING^ΔRG5^* (Martin et al., 2018); *Keap^EY5^* (Gift from Dirk Bohmann (Sykiotis and Bohmann, 2008)). BDSC: Bloomington Drosophila Stock Center, Indiana University Bloomington.

### Genotypes of clone and method clone induction

Heat shock induced somatic clones were generated using the flp/FRT technique (Xu and Rubin, 1993) or MARCM technique (Lee and Luo, 1999). Heat shock was given at 37°C for 30 minutes to early second instar progeny larvae (48 hours after egg laying, ARL) of the following parental genotypes. The larvae were then grown at 25°C till 96 to 120 hours AEL.

- ***hsrω^66^* mutant** clones (non-GFP) and its twin spot (two copies of GFP): *FRT82B hsrω^66^/ TM3b* X *y w hs-flp; FRT82B Ubi-GFP*
- **lgl^−^ clones in hsrω^66^/+ genetic background:** lgl^4^ FRT40A/Cy-O;hsrω^66^/ TM3b and *y w hs-flp tub Gal4 UAS-GFP; tub-Gal80 FRT40A/Cy-O*
- **lgl^−^ clones in Df(3R)Hrb87F)/+ genetic background:** lgl^4^ FRT40A/Cy-O;Df(3R)Hrb87F/TM3b and *y w hs-flp tub Gal4 UAS-GFP; tub-Gal80 FRT40A/Cy-O*
- ***lgl^−^ UAS-hsrω-RNAi*** and ***lgl^−^ UAS-hsrω ^EP3037^***: *lgl^4^UAS-hsrω RNAi (2X) FRT40A;* and *y w hs-flp tub Gal4 UAS-GFP; tub-Gal80 FRT40A/Cy-O.* or *lgl^4^FRT40A; hsrω^EP3037^* and *y w hs-flp tub Gal4 UAS-GFP; tub-Gal80 FRT40A/Cy-O* respectively.
- **lgl^−^ in STING^Δ5^/+ genetic background:** lgl^4^ FRT40A, STING^Δ5^/Cy-O and *y w hs-flp tub Gal4 UAS-GFP; tub-Gal80 FRT40A/Cy-O*
- ***lgl^−^ UAS-STING-RNAi:*** *w;lgl^4^ FRT40A UAS-STING RNAi/Cy-O;+* and *y w hs-flp tub Gal4 UAS-GFP; tub-Gal80 FRT40A/Cy-O*
- **lgl^−^ in Keap1^EY5^/+ genetic background:** w;lgl^4^ FRT40A Keap1^EY5^/Cy-O;+ and *y w hs-flp tub Gal4 UAS-GFP; tub-Gal80 FRT40A/Cy-O*
- ***lgl^−^ UAS-STING-RNAi:*** *w;lgl^4^ FRT40A UAS-keap1RNAi/Cy-O;+* and *y w hs-flp tub Gal4 UAS-GFP; tub-Gal80 FRT40A/Cy-O*
- ***lgl^−^ UAS-CncC:*** *w;lgl^4^ FRT40A UAS-CncC/Cy-O;+* and *y w hs-flp tub Gal4 UAS-GFP; tub-Gal80 FRT40A/Cy-O*

### Immunofluorescence staining and microscopy

Larvae of desired genotypes were dissected in 1x Phosphate Buffer Saline (PBS), and imaginal discs were fixed in 4% paraformaldehyde in 1x PBS containing 0.02 % Triton-X-100, and incubated overnight in the desired primary antibody at 4^0^C, followed by washing and incubation with appropriate fluorescently tagged secondary antibodies for two hours at the room, and counterstained with SYTOX-Green (S33025, Invitrogen) or Alexa Fluor™ 633 Phalloidin (A22284, Invitrogen). The following primary antibodies were used: anti-Caspase-3, rabbit (1:500 dilution, MERK/Sigma Aldrich, C8487), Polyubiquitinylated conjugates monoclonal antibody FK1, mouse (1:500 dilution, Enzo Life Sciences, BML-PW8805-0500), Anti-phospho Histone-3 (pSer^10^), rabbit (1:500 dilution, MERK /Sigma Aldrich H0412), anti- β Galactosidase, mouse (1:200 dilution, MERK/Sigma Aldrich, G6282). To detect MMP1, we used a cocktail of monoclonal antibodies raised in mice (3A6B4, 3B8D12, 5H7B11, and 3B8D12; 1:1:1:1 ratio at a final dilution of 1:300, Developmental Studies Hybridoma Bank (DSHB), University of Iowa), anti-Fasciclin III, Fas III, mouse, (7G10, DSHB). Anti-dSTING, rabbit (1:100, (Martin et al., 2018)). We used the following secondary antibodies from Invitrogen: Alexa Fluor™ 555 anti-mouse (A32727) and Alexa Fluor 555 anti-rabbit (A32732).

Nuclear staining was carried out using either SYTOX-Green (Invitrogen #S33025) at 1:300 or To-PRO (Invitrogen, #T3605) at 1:1000 dilution. Corticular F-Actin was labeled using Alexa Fluor™ 633 Phalloidin (#A22284, Invitrogen). After washing, the discs were mounted using a Vectashield anti-fade medium (Vector Laboratories, H-1000). Images were acquired with a Leica-SP5 confocal microscope and processed using the Leica confocal software LAS AF. Images were acquired using a 20x or a 40x (oil immersion) objective. Images of GFP-marked *lgl*; *hsrω^66^*/+ tumor-bearing larvae were acquired with a Leica M205FA stereomicroscope with epi-fluorescent illumination (excitation filter 480 nm). All images were assembled using Adobe Photoshop CS5 software.

For detecting protein aggregates, the wing imaginal discs were dissected out from larvae of the desired genotype and age in PBS and fixed in 4% formaldehyde in 1× PBS, followed by permeabilization in 0.4% Triton X-100 and 3mM ethylenediaminetetraacetic acid (pH 8.0) in 1× PBS. Fixed imaginal discs were subsequently incubated in ProteoStat detection reagent (1:10,000 dilution, Enzo Life Sciences, #ENZ-51023-KP050), washed in 1XPBS, and the wing imaginal discs were mounted in glycerol-based anti-fade and imaged using Leica-SP5 confocal microscope. We used LysoTracker® Red DND-99 dye (#L7528, Thermo-Fisher Scientific/Invitrogen) to detect acidic lysosomes and MitoTracker™ Red CMXRos (#M22425, Thermo-Fisher Scientific/Invitrogen). This cell-permeant, red-fluorescent dye stains mitochondria in live cells and is a marker for altered membrane potential. Briefly, larvae of the desired genotype were gently rinsed and dissected in 1x PBS to remove tissues like fat body, gut, etc. The flipped larvae were incubated in freshly prepared 500nM of LysoTracker (in 1X PBS) or 500nM MitoTracker (in 1X PBS) at room temperature in the dark for 15 minutes, followed by a thorough rinse in 1x PBS. Unfixed wing discs were mounted in anti-fade-containing Vectashield (Vector Laboratories, #H-1000), and images were acquired immediately using a Leica-SP5 confocal microscope and processed using Leica confocal software-LAS AF and assembled using Adobe Photoshop.

We used dihydroethidium (#D11347, Invitrogen), a free radical sensitive dye, to detect ROS levels in unfixed larval tissues (Owusu-Ansah et al., 2008). Briefly, larvae of the desired genotype were washed and dissected in 1xPBS. After removing the fat body, gut, and other adherent tissues, the flipped larvae were incubated in freshly made 1:1000 DHE at room temperature in the dark for 15 minutes. The wing imaginal discs were subsequently thoroughly rinsed in 1XPBS, mounted in anti-fade containing Vectashield (Vector Laboratories, #H-1000), and images were acquired with Leica-SP5 confocal microscope and processed using Leica confocal software-LAS AF and AdobePhotoshop CS3.

### Clone size and Fluorescence Quantification

ImageJ platform ((Schneider et al., 2012), http://rsb.info.nih.gov/ij/) was used for image analysis. For measuring size of wing discs, confocal projection images acquired with a Leica-SP5 confocal microscope were projected as a single image and used as the input file. The Freehand selection tool was used to trace the outline of the imaginal discs marked by nuclear fluorescence, and the “Measure” option was used to obtain the area of the wing disc in arbitrary units. For clone size quantification freehand selection tool was used to trace the outline of the nuclear GFP-marked clones in individual discs. For quantification of *hsrω^66^*mutant clones (marked by complete loss of GFP) and their twin spots (*hsr^+^ GFP/ hsr^+^GFP*) characterized by their brighter GFP fluorescent was calculated using “Analyze Particle” option of Image J platform on confocal projection images. Images were first converted to an 8-bit format; signal thresholds were set to calculate the number of pixels with no signal (non-GFP*, hsrω^66^*/*hsrω^66^* clones) and those with double GFP (twin spots, +/+). To quantify total fluorescence (for pH3/DHE/TRE-dsRED/γH2AX/Lysotracker/ProteoStat) confocal projection images of the red channel of individual wing imaginal discs was used as input file after converting to 8-bit formats. Image J was used to overlay a grid of uniform unit size, and the maximum fluorescence intensity per ROI was recorded. Graphs were plotted using MS Excel.

### ThT Assay for estimating protein aggregates

Larval wing discs were homogenized in ice cold 1xPBS to release cellular contents, including any insoluble amyloid-like aggregates. Thioflavin T (ThT) at a final concentration of 10μM was added to the homogenate and incubate for 10 minutes at room temperature. ThT selectively binds to β-sheet-rich protein aggregates, such as amyloids, leading to a significant increase in its fluorescence. The homogenate was transferred to a black 96-well plate and fluorescence was measured at excitation of 440–450 nm and emission at 480–490 nm using a plate reader (Bio-Rad).

### Proteomics of wing imaginal discs from *hsrω^66^* mutant and *Canton^S^* larvae

#### Protein extraction from wing discs of third instar larvae

Wing imaginal discs from third instar larvae were dissected in cold 1× PBS put in 100 μl extraction buffer (6 M GnHCl in 50 mM Tris-HCl pH 7.4, 65 mM dithiothreitol) with 50 mM sodium acetate and protease inhibitors (1× protease inhibitor cocktail with 0.2 mM PMSF).

Within 30 minutes of dissection the wing discs were sonicated with a Bioruptor (Diagenode) using 30 s ON and 30 s OFF for 5 cycles at 4°C. Cell debris was removed by centrifuging at 6000 g for 3 minutes and supernatant transferred to a new tube. The protein concentration was determined spectrophotometrically using Nanodrop. Five micrograms of the protein were used for LC-MS-MS analysis.

#### Sample preparation for LC-MS/MS

Five micrograms of the protein samples were reduced with 5 mM tris(2-carboxyethyl) phosphine (TCEP), further alkylated with 50 mM iodoacetamide and digested with Trypsin (1:50, Trypsin/lysate ratio) for 16 h at 37°C. Digests were cleaned using a C18 silica cartridge to remove the Digests were cleaned using a C18 silica cartridge to remove the salt and dried using a speed vac. The dried pellet was resuspended in buffer A (2% acetonitrile, 0.1% formic acid).

#### Mass Spectrometric Analysis of Peptide Mixtures

Experiments were performed on an Easy-nlc-1000 system coupled with an Orbitrap Exploris mass spectrometer. 1ug of peptide sample were loaded on C18 column 15 cm, 3.0μm Acclaim PepMap (Thermo Fisher Scientific) and separated with a 0–40% gradient of buffer B (80% acetonitrile, 0.1% formic acid) at a flow rate of 300 nl/min) and injected for MS analysis. LC gradients were run for 60 minutes. MS1 spectra were acquired in the Orbitrap (Max IT = 25ms, AGQ target = 300%; RF Lens = 70%; R=60K, mass range = 375−1500; Profile data). Dynamic exclusion was employed for 30s excluding all charge states for a given precursor. MS2 spectra were collected for top 12 peptides. MS2 (Max IT= 22ms, R= 15K, AGC target 200%).

#### Data Processing

All samples were processed and RAW files generated were analyzed with Proteome Discoverer (v2.5) against the Uniprot Drosophila melanogaster database. For dual Sequest and Amanda search, the precursor and fragment mass tolerances were set at 10 ppm and 0.02 Da, respectively. The protease used to generate peptides, i.e., enzyme specificity was set for trypsin/P (cleavage at the C terminus of “K/R: unless followed by “P”). Carbamidomethyl on cysteine as fixed modification and oxidation of methionine and N-terminal acetylation were considered as variable modifications for database search. Both peptide spectrum match and protein false discovery rate were set to 0.01 FDR.

#### Proteome data analysis

To identify biologically relevant protein signatures in *hsrω^66^* wing disc proteome versus wild type proteome we calculated the log2 abundance ratios, using mean abundance values for individual UniProt IDs. Only those with combined FDR confidence <0.05 (medium) or <0.01 (high) were taken into consideration; those with combined FDR >0.05 (low) were discarded. We further filtered out peptides that were not detected in either MS or MS/MS spectra depending on the peak calling. Overall 3125 Uniprot IDs were subjected to function enrichment analysis using STRING platform (https://string-db.org/, (Szklarczyk et al., 2021)) in conjunction with ssGSEA (single-sample Gene Set Enrichment Analysis).

### Kaplan-Meier survival plots

Kaplan-Meier survival plots were generated using clinical data from TCGA (The cancer genome atlas), using cBioPortal platform (https://www.cbioportal.org/). Datasets were restricted to epithelial cancers only and queried for CNA (Copy Number Alterations) and mutations for individual genes: *KEAP1*, *HSF1* and *STING1*.

### Study of *Sat III* germline copy number variations in PCa patients

#### Study population and patient recruitment

The peripheral blood used in this study was collected from the biopsy-confirmed treatment-naïve prostate cancer (PCa) patients (n=18) and benign prostatic hyperplasia (BPH) patients (n=5), enrolled at the Department of Urology, King George’s Medical University (KGMU, Lucknow, India). Clinical specimens were collected after obtaining written informed consent and institutional review board approvals from KGMU and Indian Institute of Technology Kanpur (Kanpur, India). The study was initiated after obtaining approval from the Institutional Ethics Committee.

### Digital droplet PCR quantification of SatIII copy number in PCa patients

Blood was collected in K_2_-EDTA 2ml tubes from BPH (benign prostatic hyperplasia) and PCa (prostate cancer) patients. Genomic DNA was isolated using QIAamp® DNA Blood mini-kit (#51104, Qiagen), which was further processed to identify *Sat III* copies. Droplet digital PCR was performed with the QX200 Droplet Digital PCR System (#1864001,Bio-Rad, USA). The ddPCR reaction mixture consisted of 10μL of 2XddPCR Supermix for Probes (#1863026, Bio-Rad, USA), 10pg of genomic DNA, and 20Xprimers/probes mix in a final volume of 20μL. The entire reaction mixture was loaded into a sample well of a DG8 Cartridge (Bio-Rad, USA) together with 70μL of droplet generation oil (Bio-Rad, USA) and placed into the automated droplet generator (Bio-Rad, USA). After processing, the droplets generated from each sample were transferred to a 96-well PCR plate (Bio-Rad,USA). PCR amplification was carried out on a T100 Touch thermal cycler (Bio-Rad, USA) using a thermal profile beginning at 95 °C for 10 min, followed by 40 cycles of 94 °C for 30 s, and 60 °C for 60 s, and ending of 98 °C for 10 min at a ramp rate of 2 °C/s. After PCR, the plate was loaded on the droplet reader (Bio-Rad, USA); acquired data were analyzed with QuantaSoft Analysis Pro software (Bio-Rad, USA). All samples had ≥10000 accepted droplets to be considered for analysis. SatIII copies were determined by QuantaSoft, the software coupled to the Droplet Reader QX200 that applies a Poisson algorithm to calculate the initial concentration of DNA target molecules as units of copies/μL input based on positive and negative droplets. Specifically, after setting an appropriate fluorescence threshold to determine the number of positive droplets (containing at least one target molecule), the software calculates the number of copies per μL of the reaction volume.

Student’s *t*-test was applied to determine the statistical significance for independent samples p-value. Violin plots show the median and quartiles.

### Sat III expression analysis in cancer cell lines

#### Cancer cell lines and culture conditions

22RV1, PC3, VCaP, LNCaP (Prostate cancer cell lines), were obtained from American Type Culture Collection (ATCC) and were cultured as per the ATCC recommended guidelines. Briefly, the cells were cultured in a growth medium supplemented with 10% fetal bovine serum (FBS) and 0.5% Penicillin-Streptomycin (Gibco, Thermo-Fisher Scientific) in a cell culture incubator (Thermo Fisher Scientific) supplied with 5% CO_2_ at 37°C.

#### RNA isolation and Quantitative PCR (qPCR)

The total RNA was extracted using RNAiso Plus (Takara), and 1µg of total RNA was reverse transcribed into cDNA using First Strand cDNA synthesis kit (PureGene) according to the manufacturer’s protocol. Quantitative PCR (qPCR) was performed using cDNA template, primer sets (sat III_FP: TATGAATTCAATCAACCCGAGTGCAATCGAA, sat III_RP: TATGGATCCTTCCATTCCATTCCTGTACTCG; GAPDH_FP: TGCACCACCAACTGCTTAGC, GAPDH_RP: GGCATGGACTGTGGTCATGAG) and SYBR Green PCR Master-Mix (PureGene) on the QuantStudio 5 Real-Time PCR System (Applied Biosystems). Relative sat III expression was calculated for each sample by the ΔΔC_T_ method. A two-tailed unpaired Student’s t-test was applied to determine the statistical significance for independent samples *P≤ 0.05 and **P≤ 0.001. Error bars represent mean ± SEM.

## Supplementary data

**Figure S1 Comparative Proteome of wing epithelium of *hsrω^66^* mutants versus wild type**

**(A-B)** Box plot displaying abundance of proteins in *hsrω^66^* mutant and wild type control proteomes (A); Volcano plot of 3198 variables depicting the comparative levels of mis-regulated proteins in *hsrω^66^* proteome compared to wild type (B).

**(C-D)** Relative abundance (log_2_ fold change) of select proteins belonging to Protein metabolism (C) and Stress response (D) functional classes in *hsrω^66^* proteome compared to its wild type counterpart.

**(E)** Rank-based Gene Ontology (GO) enrichment analysis of RNA polII transcription regulatory complex (GO: 00905757) and Mediator complex. The x-axis represents the list of proteins in ascending order of fold change in *hsrω^66^* versus wild type wing disc proteome. Black bars along the x-axis mark the position of proteins being queried. The tables alongside give the protein levels of the top five mis-expressed proteins belonging to a given GO class.

(**F**) Representative examples of wing imaginal discs displaying increase in ubiquitinated proteins using anti-Ubiquitin FK2 Antibody in *hsrω^66^* and *hsrω^66^*/+ wing imaginal discs.

**Figure S2. *lgl* clones display cellular stress**

**(A-C)** GFP-marked *lgl* mutant cells generated in a wild type epithelia exhibit signatures of cellular stress. *lgl* clones (GFP, green) display elevated levels of ROS, (DHE, Red, A), increase in MitoTracker staining, revealing perturbed mitochondrial activity (B), and elevated number of acidic lysosomes as seen by LysoTracker staining (red, C). Area marked by yellow box magnified on the right in each case. Scale bars: 100 µm.

**Figure S3. Genetic disruption of omega biocondensates facilitates *lgl* tumor progression (A-B)** GFP-marked *lgl* clones generated in *Df(3R)Hrb87F/+* genetic background (*lgl;Df(3R)Hrb87F/+,* A) or with co-downregulation of ISWI (*lgl;UAS-ISWI-RNAi,*) undergo neoplastic transformation, as revealed by disrupted F-Actin. Transformed clones exhibit loss of junction protein FasIII (**A**). Scale bars: 100 µm.

**Figure S4 *hsrω*/+ host genetic background facilitates growth of *lgl* tumors in *Drosophila* adult gut**

(**A-C**) GFP-marked somatic control (wild type) (A) or with loss of *lgl* (B, C) clones generated in wild type (A, B) or *hsrω^66^/+* (C) genetic background. Note significantly large *lgl* clones in C compared to A or B, with characteristic small nuclei (red, stained with To-PRO) indicative of intestinal stem cells.

**Figure S5. Cellular Localization of STING**

**(A)** Uniform punctae of dSTING in cells of larval salivary gland over expressing dSTING (*vg Gal4 >UAS-dSTING*).

**Figure S6. Genetic variations in *Sat III* in human cancer patients**

**(A-B)** Kaplan-Meier survival curves generated on cBioPortal (cbioportal.org): comparing survival of individuals with epithelial cancer carrying Copy Number Alterations (CNA) in *KEAP1* with individuals not carrying an alteration in *KEAP1*. The plot was generated by querying 91469 samples across 233 studies, across 16 cancer types (A). The inset displays distribution of various genetic alterations in *HSF1* across patients (B); comparing survival of individuals with cancers of Lymphoid origin, carrying Copy Number Alterations (CNA) in *STING1*. The plot was generated by querying 9672 samples across 17 studies.

## Notes

### Competing Interest Statement

The authors have declared no competing interest.

### Summary of Updates

The revised version of the manuscript carries the following addition: 1.Proteomics: Proteome analysis of the lncRNA hsr-omega mutant and control larval epithelium that provides mechanistic insights into the role of hsr-omega in regulating protein homeostasis or proteostasis. Corresponding to Figure 2. 2.Inclusion of two new genetic regulators of cancer susceptibility, namely the innate immune cGAS-STING pathway member dSTING; and the redox regulators Nrf2 and Keap1. (New Figure 4.) 3.Chemical rescue of Drosophila tumor phenotypes by use of Proteosome inhibitor and STING agonist. (addition to Figure 3 and Figure 4) 4.Inclusion of germline copy number variations in Satellite III in Prostate cancer patients. (Revised Figure 5)

