## Supplemental Figure Legends and Figures for "Nuclear biocondensate-forming lncRNA and cellular stress surveillance shape host cancer susceptibility"

**Figure S5. Cellular Localization of STING**

**(A)** Uniform punctae of dSTING in cells of larval salivary gland over expressing dSTING (*vg Gal4 >UAS-dSTING*).

**Figure S6. Genetic variations in *Sat III* in human cancer patients**

**(A-B)** Kaplan-Meier survival curves generated on cBioPortal (cbioportal.org): comparing survival of individuals with epithelial cancer carrying Copy Number Alterations (CNA) in *KEAP1* with individuals not carrying an alteration in *KEAP1*. The plot was generated by querying 91469 samples across 233 studies, across 16 cancer types (A). The inset displays distribution of various genetic alterations in *HSF1* across patients (B); comparing survival of individuals with cancers of Lymphoid origin, carrying Copy Number Alterations (CNA) in *STING1*. The plot was generated by querying 9672 samples across 17 studies.

Figure S1

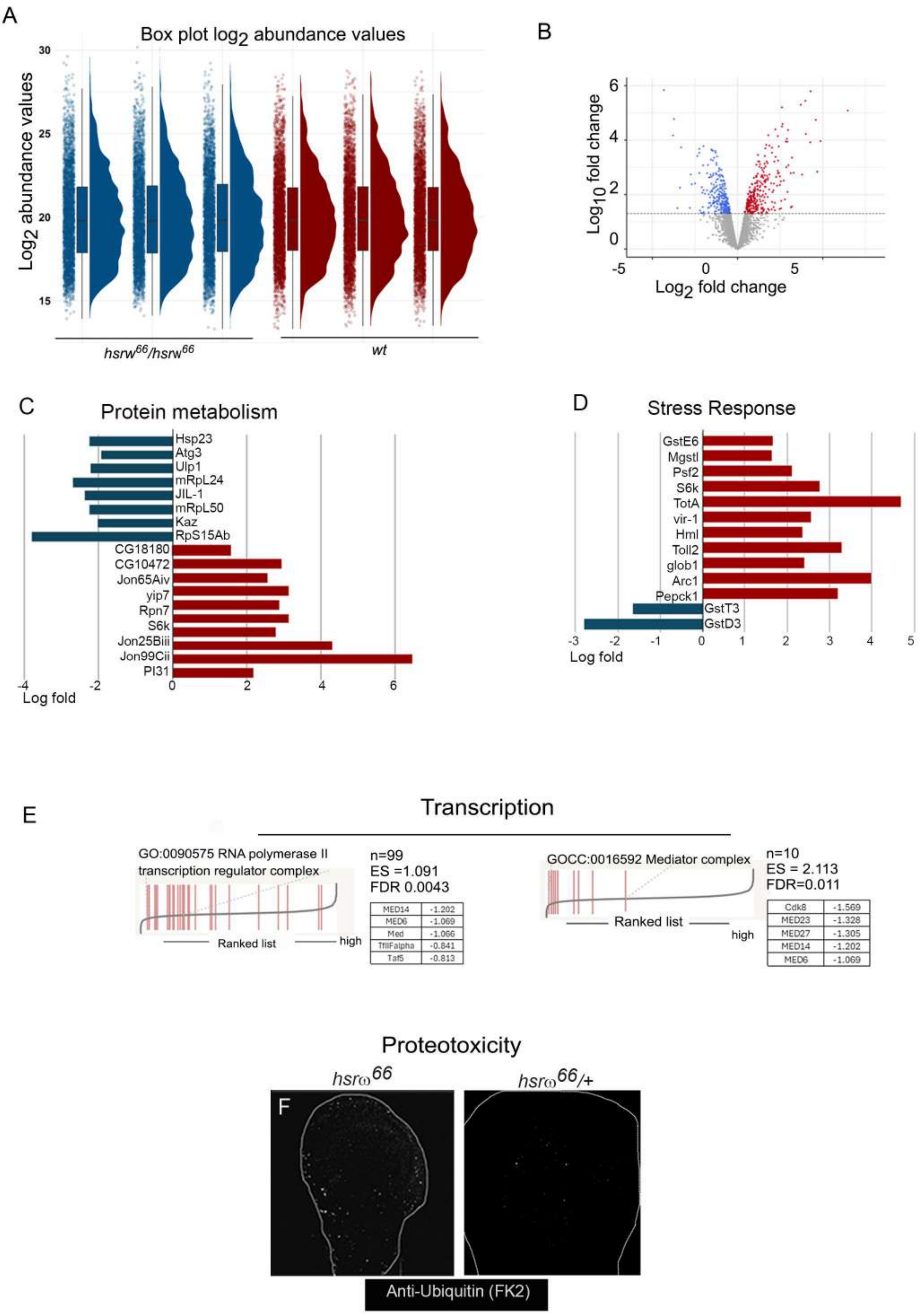
A
Box plot log2 abundance values
Log2 abundance values
30
25
20
15
hsrw66/hsrw66
wt
B
Log10 fold change
Log2 fold change
6
4
2
0
-5
0
5
C
Protein metabolism
Hsp23
Atg3
Ulp1
mRpL24
JIL-1
mRpL50
Kaz
RpS15Ab
CG18180
CG10472
Jon65Aiv
yip7
Rpn7
S6k
Jon25Biii
Jon99Cii
Pl31
-4
-2
0
2
4
6
Log fold
D
Stress Response
GstE6
Mgstl
Psf2
S6k
TotA
vir-1
Hml
Toll2
glob1
Arc1
Pepck1
GstT3
GstD3
-3
-2
-1
0
1
2
3
4
5
Log fold
E
Transcription
GO:0090575 RNA polymerase II transcription regulator complex
Ranked list
high
n=99
ES =1.091
FDR 0.0043
MED14 -1.202
MED6 -1.069
Med -1.066
TfIIFalpha -0.841
Taf5 -0.813
GOCC:0016592 Mediator complex
Ranked list
high
n=10
ES = 2.113
FDR=0.011
Cdk8 -1.569
MED23 -1.328
MED27 -1.305
MED14 -1.202
MED6 -1.069
Proteotoxicity
hsrω66
hsrω66/+
F
Anti-Ubiquitin (FK2)

Figure S2

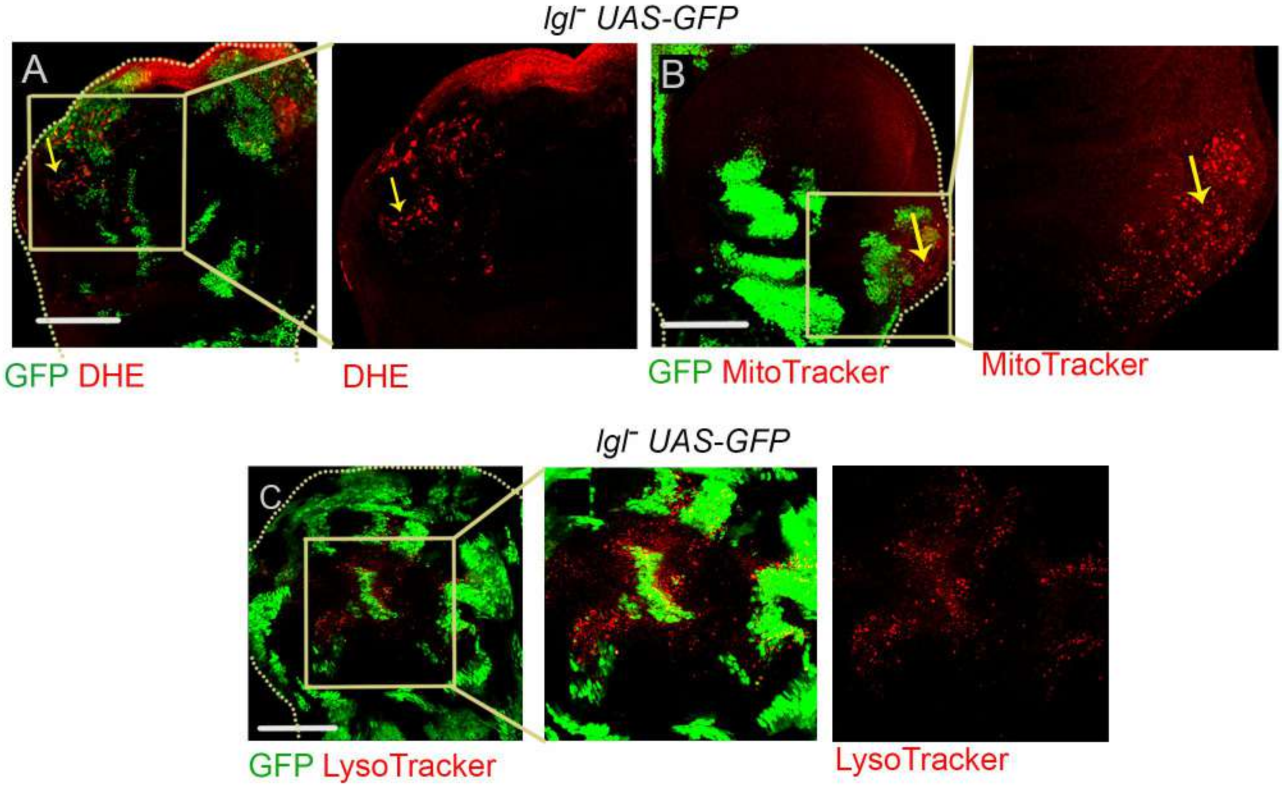

lgl⁻ UAS-GFP
A
GFP DHE
DHE
B
GFP MitoTracker
MitoTracker
lgl⁻ UAS-GFP
C
GFP LysoTracker
LysoTracker

### Figure S3

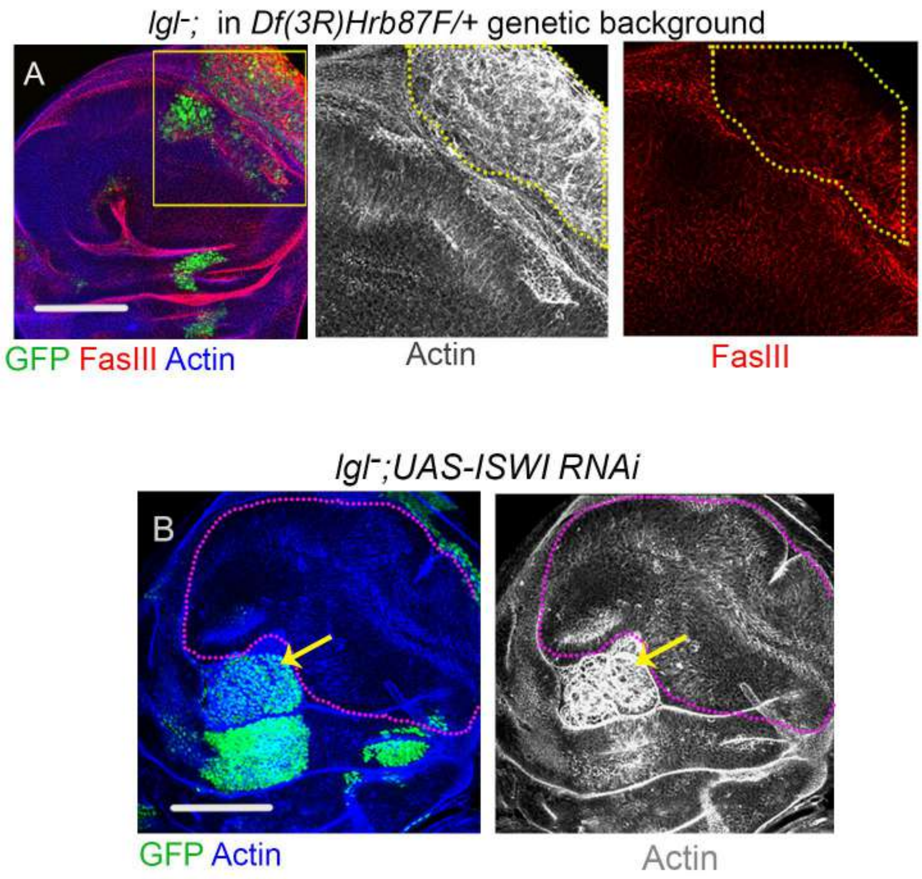

lgl-; in Df(3R)Hrb87F/+ genetic background
A
GFP FasIII Actin
Actin
FasIII
lgl-;UAS-ISWI RNAi
B
GFP Actin
Actin

Figure S4

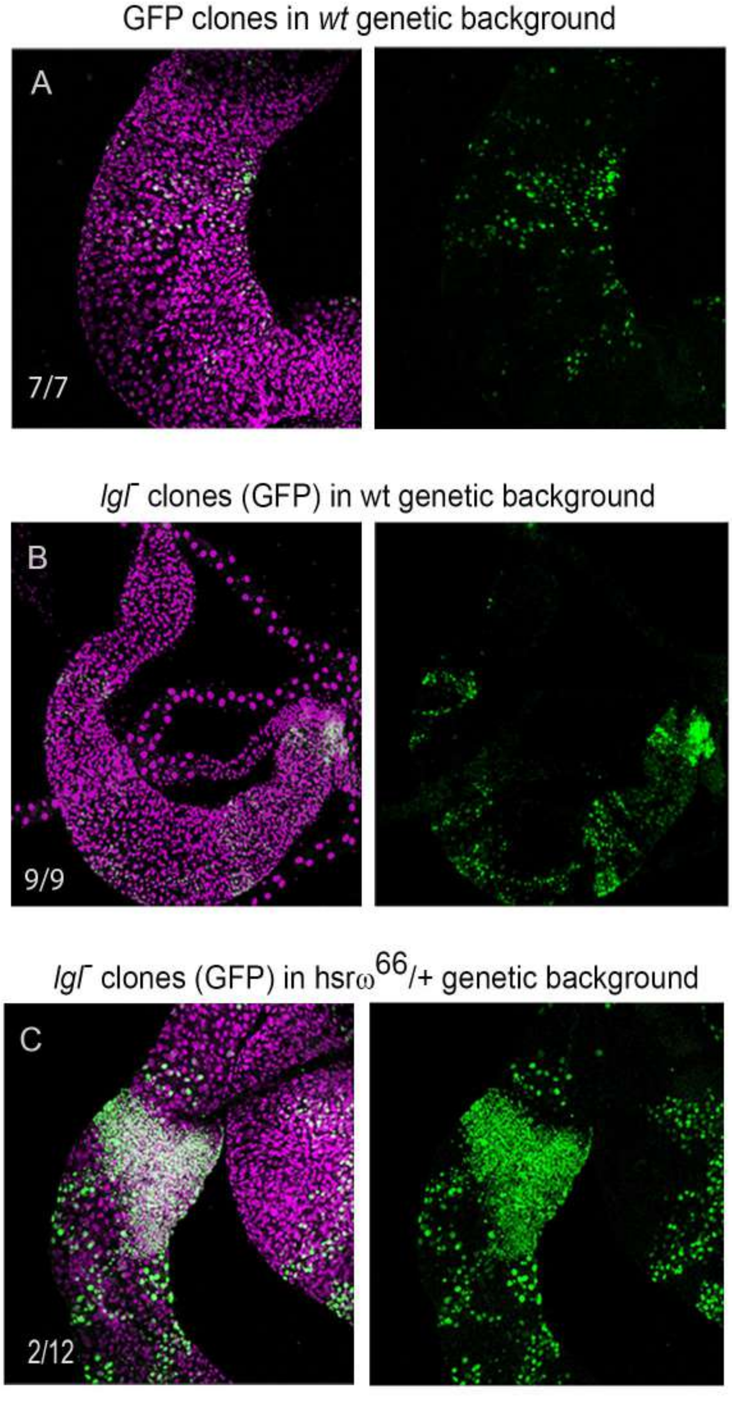
GFP clones in wt genetic background
A
7/7
lgl- clones (GFP) in wt genetic background
B
9/9
lgl- clones (GFP) in hsrω66/+ genetic background
C
2/12

Figure S5

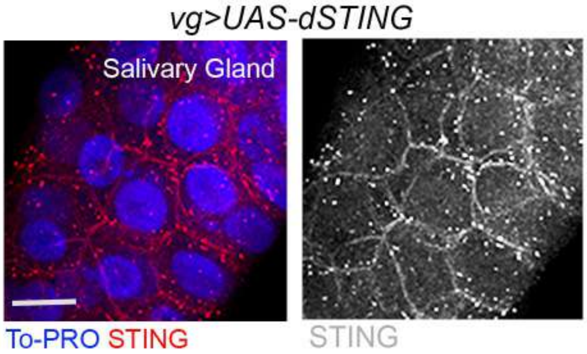

Figure S6

A

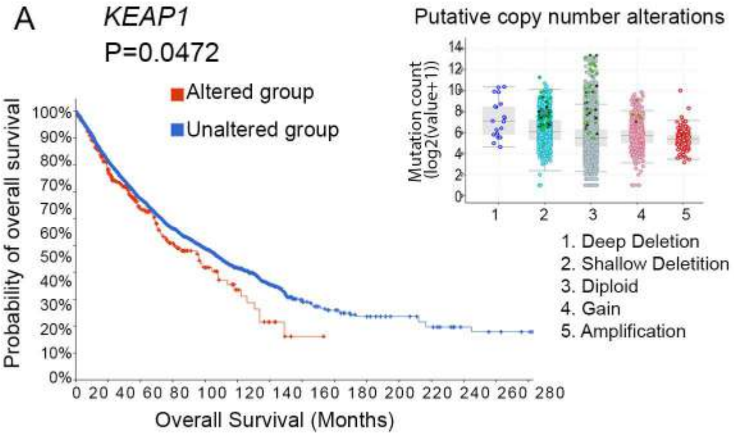
KEAP1
P=0.0472
Altered group
Unaltered group
Probability of overall survival
100%
90%
80%
70%
70%
60%
50%
40%
30%
20%
10%
0%
0 20 40 60 80 100 120 140 160 180 200 220 240 260 280
Overall Survival (Months)
Putative copy number alterations
Mutation count (log2(value+1))
14
12
10
8
6
4
2
0
1 2 3 4 5
1. Deep Deletion
2. Shallow Deletition
3. Diploid
4. Gain
5. Amplification

B

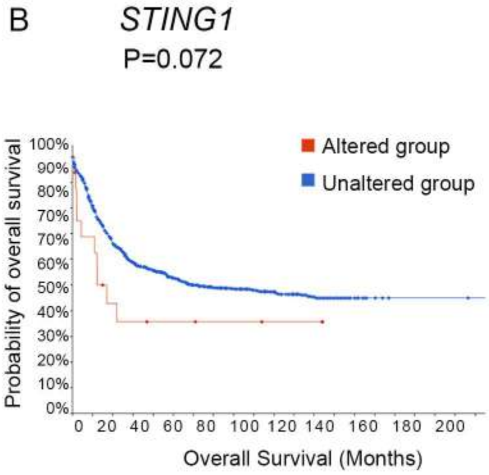
STING1
P=0.072
Altered group
Unaltered group
Probability of overall survival
100%
90%
80%
70%
70%
60%
50%
40%
30%
20%
10%
0%
0 20 40 60 80 100 120 140 160 180 200
Overall Survival (Months)
